# Convolutional and Recurrent Neural Network for Human Action Recognition: application on American Sign Language

**DOI:** 10.1101/535492

**Authors:** Hernandez Vincent, Suzuki Tomoya, Venture Gentiane

## Abstract

Human Action Recognition (HAR) is an important and difficult topic because of the important variability between tasks repeated several times by a subject and between subjects. This work is motivated by providing time-series signal classification and a robust validation and test approaches. This study proposes to classify 60 American Sign Language signs from data provided by the LeapMotion sensor by using a combined approach with Convolutional Neural Network (ConvNet) and Recurrent Neural Network with Long-Short Term Memory cells (LSTM) called ConvNet-LSTM. Moreover, a complete kinematic model of the right and left forearm/hand/fingers/thumb is proposed as well as the use of a simple data augmentation technique to improve the generalization of neural networks. Results showed an accuracy of 89.3% on a user-independent test set with data augmentation when using the ConvNet-LSTM, while LSTM alone provided an accuracy of 85.0% on the same test set. The result dropped respectively to 85.9% and 81.4% without data augmentation.

## Introduction

Sign language is a language that mainly uses hand kinematics and facial expressions. It is widely used by hearing-impaired people to communicate with each other, but rarely with people who do not have hearing impairments. Therefore, they are only in direct contact with hearing-impaired persons, which greatly limits social interactions. An alternative would be to have a real time translation with interpreters but they cannot be permanently available and can be quite expensive. Therefore, a system that could enable automatic translation would be of great interest.

Human Action Recognition (HAR) in general is an important and challenging topic because of the large variability that exists for a given task. Indeed, whether the variability comes from a subject that repeat an action several times or more importantly between subjects, the kinematics behaviour over time presents a certain challenge to generalize. HAR considers that this dynamics behaviour may be represented by specific patterns that could be classified using machine learning algorithms.

Nowadays, human movement data can be easily extracted from low-cost systems that integrate depth mapping sensor such as Kinect and LeapMotion. These systems are ready to use, require a relatively short time to set up and the data is easy to extract. Therefore, they can be easily used to quickly acquire a large amount of data, which is a requirement when considering machine learning. Other approaches have been considered with sEMG (Savur and Sahin 2016), CyberGloves (Sarawate et al. 2015) or motion capture system but these techniques are difficult to use outside of laboratory.

Considering systems such as the Kinect or a combined Kinect/LeapMotion approach, several studies have been conducted using video or depth-mapping sensor and machine learning approaches such as Hidden-Markov-Model (HMM) (Zafrulla et al. 2011), coupled HMM (Kumar et al. 2017), random-forest per-pixel (Dong et al. 2015), multi-class SVM (Marin et al. 2014), linear binary SVM (Weerasekera et al. 2013), ConvNet-LSTM video based approaches (Yang and Zhu 2017) to name a few. Such approaches are interesting but the Kinect remains difficult to use in public space since it requires large space and a power supply.

The LeapMotion easily detects forearm and palm movements as well as fingers and thumb joints position. In addition, it can be used with a single USB port and allows a measurement precision of 200 μm (Weichert et al. 2013) to estimate the joint position. Therefore, it seems to be an interesting choice as a basis for sign language recognition and for the development of future portable systems. Naidu and Ghotkar (2016) used the depth-mapping from the leap motion to classify a subset of the Indian Sign Language (10 Arabic numbers, 26 letters and 9 words). They propose four different approaches (Euclidian distance measure, similarity, Jaccard and dice similarity) with 8 distances computed from 6 measured points (centre of the palm and all finger tips). The cosine similarity showed an accuracy of 90.00% for the complete dataset but their approach is limited to static posture. Marin, Dominio and Zanuttigh (2014) studied a combined LeapMotion/Kinect approach with multi-class SVM to classify 10 static words from the ASL with key-moment manually extracted. They presented an average accuracy of 80.86%, 89.71% and 91.28% with the LeapMotion, the Kinect and combined Kinect/LeapMotion approaches respectively with a user independent K-fold cross-validation (M-1 subject for each K) for parameters tuning. Then, the classifier was trained again with all subjects (M) and computed the accuracy on it which does not provide a model that can be used with new subjects. Kumar, Gauba, Roy and Dogra (2017) also considered a combined Kinect/LeapMotion approach on 25 dynamics words of the Indian Sign Language. They considered a coupled-HMM approach and gained a 90.8% of accuracy on 25% out of the complete dataset (50% used for training and 25% for parameters tuning and parameters validation), however they do not provide information whether a user-independent test was considered or not. Chuan et al. (2014) classified the 26 letters of the English alphabet in the American Sign Language by using k-NN and SVM classifier. Results showed an average classification rate of 72.78% and 79.83% respectively using K-fold cross-validation (K=4) on the complete dataset composed of 2 subjects. Fok et al. (2015) proposed an HMM based approach to recognize the 10 ASL Arabic numbers with an overall average recognition of 93.14% by using for each subject half of the sample for training and the rest to test the system.

The recent breakthrough of deep learning has outperformed conventional machine learning approach in computer vision task (Lee et al. 2009). Deep learning uses a succession of multiple layers to extract information, with the output of one layer as the input of the next one. They provide a robust approach to generalize inter-subject variability and they can take into account the time-series dynamics behaviour of human movements mostly with convolutional (ConvNet) and Recurrent Neural Network (RNN). Regarding the sign language recognition, Koller et al. (2016) use an hybrid ConvNet-HMM approach based on sequence of images extracted from video and by tracking the dominant hand.

Regarding deep learning, the state of the art on HAR is presented by Ordóñez and Roggen (2016) adapted from Sainath et al. (2015) for speech recognition. They proposed an approach based on combining a ConvNet and a RNN with Long-Short-Term-Memory (LSTM) for HAR with data from Inertial Measurement Unit sensors (IMUs). ConvNet and LSTM are neural networks with a supervised learning approach that can learn the dependencies between some given inputs and outputs. ConvNet and LSTM combined outperformed both approaches considered independently for speech recognition (Sainath, Vinyals, Senior and Sak 2015) and HAR (Ordóñez and Roggen 2016). Indeed, both approaches have their advantages with ConvNet that can extract features from a given signal and LSTM that can consider the dynamics in a time-series signal (Gers et al. 2000, Hochreiter and Schmidhuber 1997, Lipton et al. 2015). A combined approach seems to be the best compromise to perform HAR and is promising since it allows flexible data fusion and features extraction (Ordóñez and Roggen 2016).

To the best of our knowledge, no studies that used the LeapMotion for sign language recognition performed a robust user independent K-fold cross-validation and test. Indeed, deep learning model can easily learn the behaviour of subjects but failed on new subjects. This research proposes an approach based on a state of the art combination of a convolutional neural network (ConvNet) to extract relevant features from a complete right and left forearm/hand/fingers/thumb kinematics model followed by a Recurrent Neural Network with long-short term memory cells (LSTM) to classify 60 signs of ASL. Kinematics is derived from the skeletal tracking model provided by the Leap Motion sensor. The emphasis of the research is also on using a robust training and validation approach that corresponds to an independent K-fold cross-validation for hyperparameters tuning, followed by a user-independent test. It is also demonstrated that training and testing approach that is not user-independent can lead to overestimated accuracy.

## Material and methods

### Experimentation

A dataset of 25 male subjects, all novices in any sign language, was collected. Before each measurement, the corresponding sign was taught to them. In this study, the Arabic numbers from 0 to 10 and 49 words were considered. The total numbers of each sign gathered are presented in the Table 1.

**Table 1.**
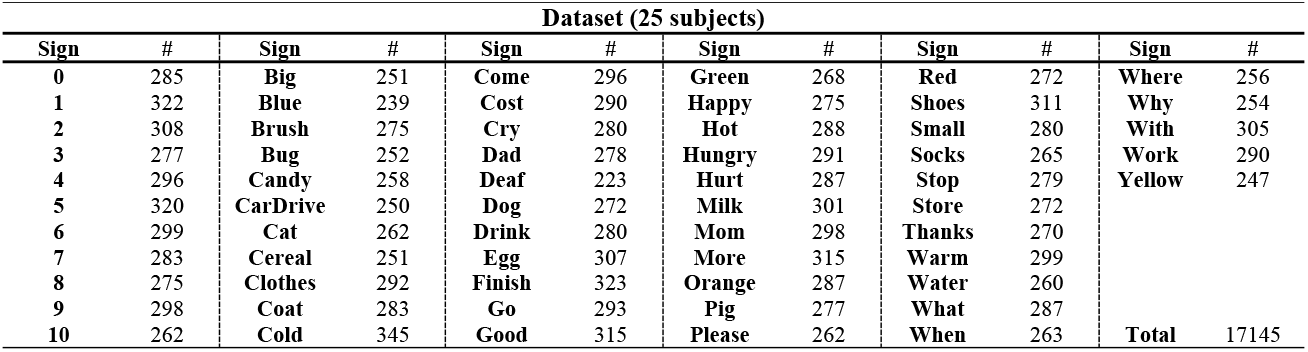
Numbers of sign gathered during the experiment with the 25 subjects.

### Features extraction

The LeapMotion SDK Unity core assets 4.3.2 was used with the C#/Unity module to obtain the data during the experimental protocol. A total numbers of 26 points on each hand can be extracted in real-time from the LeapMotion to represent the kinematics of the finger/hand/forearm system. The features computed from these points are inspired from a multi-finger modelling approach presented by Carpinella et al. (2011) with minor modifications and the ISB recommendations for the relative orientation of the hand to the forearm (Wu et al. 2005).

### Kinematics models

The following coordinates systems are considered for the hand (1) and forearm (2) coordinates system (Figure 1). (1) 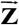 has the direction of (M_*2*_-M_*5*_) pointing externally, 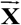 is perpendicular to the plan form by (M_*2*_-RS) and (P_*5*_-RS) pointing forwardly, and 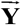 is the cross product of 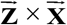 pointing upwardly. (2) 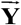 has the direction of (EL-US) pointing upwardly, 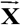 is the cross product of 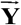 and (RS-US) pointing forwardly and 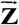 is the cross product of 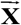 and 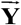 pointing externally. Then, the Euler angles that describe the relative orientation of the hand with the forearm (flexion/extension and radial/ulnar deviation) are computed with a **ZXY** rotation sequence. All vector direction are expressed from the standard anatomical position.

**Fig. 1.**
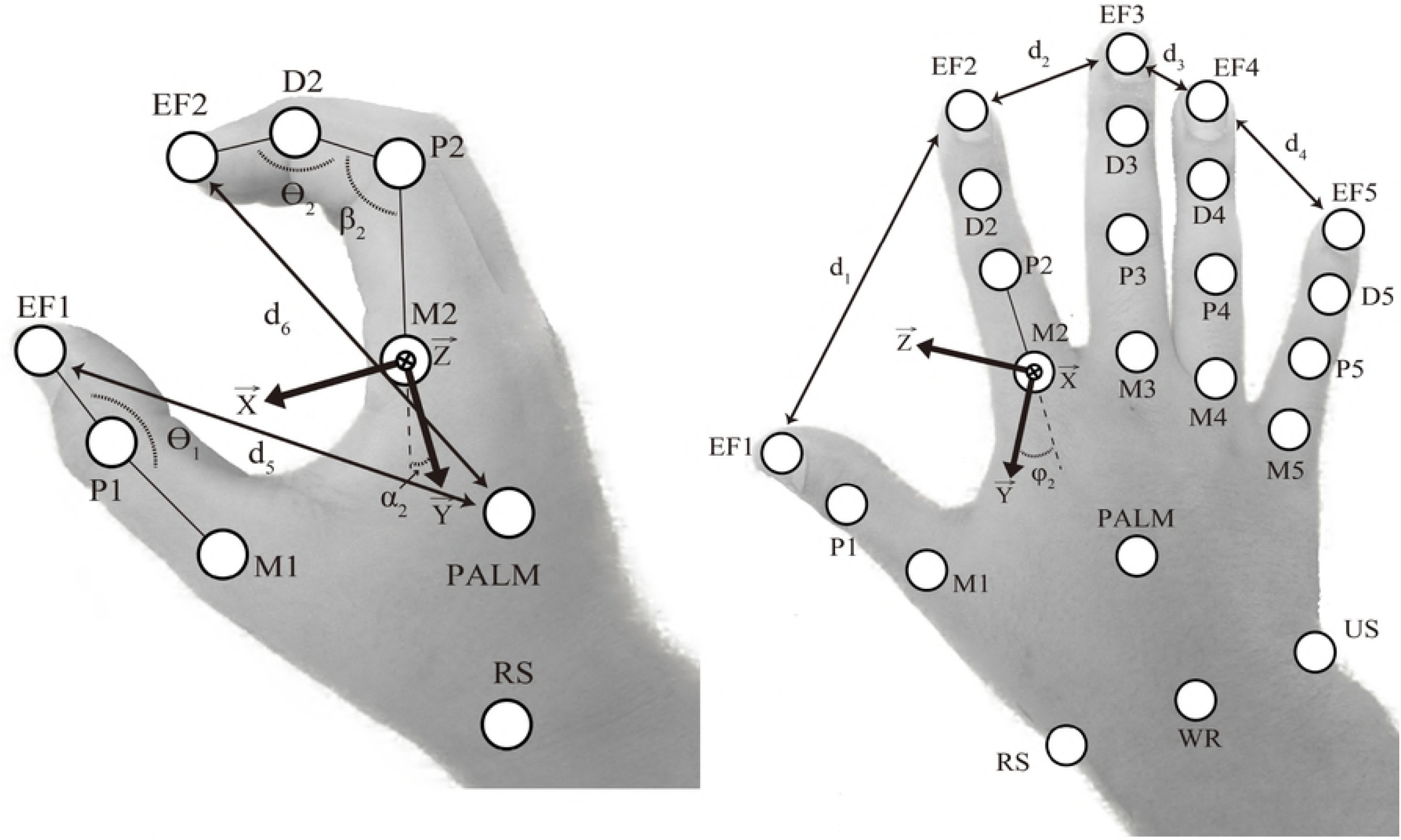
Hand joints. Thumb, index, middle, ring and pinky are numbered from 1 to 5 respectively. EF_*i*_ (*i*=1-5): finger and thumb end-effector position, M_*i*_ (*i*=1-5): finger and thumb metacarpophalangeal joint, P_*i*_ (*i*=1-5): finger and thumb proximal interphalangeal joints, D_*i*_ (*i*=2-5): finger distal interphalangeal joints. EL: elbow joint (not represented here), WR: wrist joint, RS: radial styloid, US: ulnar styloid, PALM: palm center.

The following kinematics are considered for the thumb and fingers (thumb: *i*=1, index: *i*=2, middle: *i*=3, ring: *i*=4 and pinky: *i*=5) with EF_*i*_ (*i*=1-5) that represents the finger and thumb end-effector, M_*i*_ (*i*=1-5) the finger and thumb metacarpophalangeal joint, P_*i*_ (*i*=1-5): the finger and thumb proximal interphalangeal joints and D_*i*_ (*i*=2-5) the finger distal interphalangeal joints. Then, the relative orientation of each finger with the hand is represented with a flexion angle α_i_ (Angle between 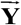 and M_*i*_ - P_i_ projected in the **XY** plan) and an abduction angle φ_i_ (Angle between 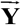 and M_*i*_ - P_i_ projected in the **YZ** plan). Moreover, the relative orientation of the thumb with the hand is represented with a flexion angle α_i_ (Angle between 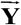 and M_*i*_ - D_i_ projected in the **YZ** plan) and an abduction angle φ_1_ (Angle between 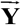 and M_*i*_ - D_i_ projected in the **XY** plan). Furthermore, the angle β_i_ between P_*i*_ - D_*i*_ and P_*i*_ - M_i_ and the angle ϴ_i_ between EF_i_ - D_*i*_ and P_i_ - D_*i*_ for the *i^th^* finger (*i*=2-5) and the angle ϴ_1_ between EF_1_ - D_1_ and M_1_ - D_1_ for the thumb are also computed. Finally, d_*j*_ (*j*=1-9) represent respectively the Euclidian distance between each successive EF (EF_1_ with EF_2_, EF_2_ with EF_3_ …) and between EF_*i*_(*i*=1-5) and the Palm.

### Dataset

Each sign is manually extracted and interpolated to 100 frames with a cubic spline interpolation. The dataset was composed of a total number of *d* = 17145 labelled signs. *f* = 60 features are computed to represent the kinematics of the right and left sides (Figure 1). For each sign extracted, the features were regrouped in a time-series matrix **F**_*l*_ ∈ ▯^*m*×*n*^ (*m* = 100 and *n* = 60) with each group given by: (1) the finger and thumb end-effector (EF) relative distance, (2) their distance with the palm, (3) the relative orientation of the hand with the forearm, (4) the relative orientation of the fingers and thumb with the hand and (5) the thumb and finger angles.

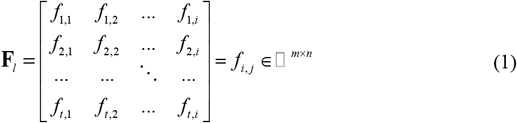

All feature matrices were computed for all signs (*l*) and subjects (*k*). Finally, each feature *f_i,j_* was standardized with the mean *μ* and standard *σ* deviation of their respective group presented above as follow:

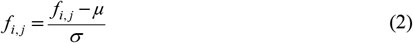

Then, each subject in the dataset is represented as 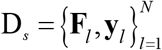 with *N* the numbers of sign and **y**_*l*_ the corresponding outputs labels of **F**_*l*_ represented as a binary vector.

### Data augmentation

To prevent the overfitting of the model and help generalize the classification, data augmentation was considered. A given signal **f**_*i*_ is sliced into 10 parts of equal length. Then, data warping was considered on the magnitude and the temporal location of the signal that were modified as follows. First, each part was distorted to a random value *p* {*p* ∈ ▯ |5 ≤ *i* ≤ 15} with a cubic spline interpolation and all parts were reconstructed and interpolated to 100 frames. Then, the amplitude of the signal was slightly altered. A sinus wave with a random period *P* {*P* ∈ ▯ |0.5 ≤ *P* ≤ 2}, phase φ {*φ* ∈ ▯ |0 ≤ *φ* ≤ *π*} and amplitude *A* {*A* ∈ ▯ | −0.2 ≤ *A* ≤ 0.2} was generated. Finally, this sinus wave was multiplied by *A* and by the amplitude range of the current signal **f**_*i*_ and added to it, thus allowing a smooth variation. This procedure is repeated 40 times for each **F**_*l*_ in the training and validation set (described in the next section).

## Gesture classification

### Training, validation and test set

The dataset was separated into a training/validation set and a test set composed respectively of 20 and 5 subjects randomly selected. The test set was kept aside to assess the final performance of the selected models. Then, a non-exhaustive user independent K-Fold cross-validation was performed on the training/validation set tha consisted of a rotating *K* = 4 folds with 15 and 5 subjects. This procedure was used to stop the training when the accuracy of the test set increases but the accuracy of the validation set decreases (overfitting of the model on the training set). Moreover, it was also used to tune the hyperparameters of the classifier (e.g. number of layers, cells unit size, learning rate, strides, patch size…). Following this procedure, the final performance of the selected classifier was estimated with the test set. The final numbers of labelled data is 542,640 for the train/validation set (with data augmentation) and 3,579 for the test set (without data augmentation).

### Training properties

The ConvNet-LSTM was trained with mini-batches composed of 500 samples and a learning rate of η = 0.001 with an exponential decay of 0.9 every 5 epochs. As a form of regularization, a dropout wrapper is added on each layer to randomly select units that are ignored at each epoch with a probability value of 0.8. The Adam gradient descent optimization algorithm (Kingma and Ba 2014) was used to minimize the cost function *E* that corresponds to a softmax cross-entropy between the estimated vector (logits) (**y’**) and the true labels vector (**y**):

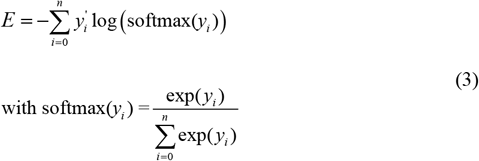

For each K-Fold cross-validation, the training phase was stopped when the accuracy of the validation set start to decrease and the corresponding ConvNet-LSTM was saved.

### ConvNet-LSTM hyperparameters

The considered hyperparameters were a ConvNet composed of 4 convolutionnal layers used to extract revelant feature in the input signal (Figure 2). Each layer receives the raw signal (or the one from the previous layer) and performs a convolution throught a 1 dimensional kernel without pooling on each **f**_*i*_ (columns) to extract revelant patterns (Ordóñez and Roggen 2016). Then, the ouput of the ConvNet is fed into 2 LSTM layers composed of the LSTM cell from Zaremba et al. (2014) with an hidden state size of 200. Tensorflow 1.4 (Mart et al. 2016) with Python 3.5.2 was used and the training was performed on a GTX 1060 6GB that integrate 1280 CUDA cores (Nickolls et al. 2008).

**Fig. 2.**
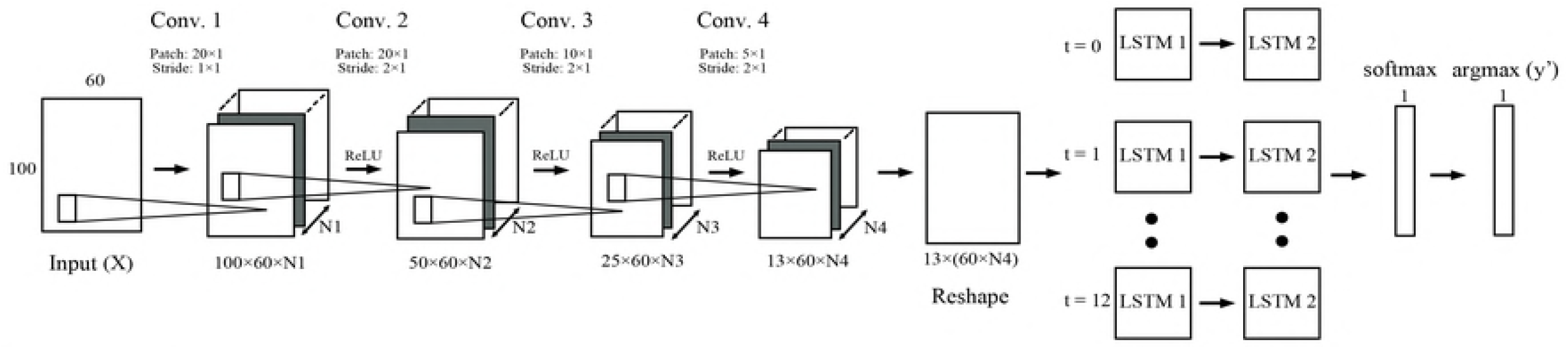
ConvNet-LSTM architecture. N1, N2, N3 and N4 represents the numbers of channel (also called feature map) created after each convolutional layer. Their values are respectively 16, 16, 32 and 64. ReLU (rectified linear unit) represent the activation function.

### LSTM hyperparameters

To show the improvement gained with the combined approach with a ConvNet and a LSTM, the same training parameters described in the section 3.b and 3.b were used to train a LSTM neural network composed of 4 LSTM layers with the same LSTM cells used in the ConvNet-LSTM with a hidden state size of 300. The LSTM-part in the two models was different since the hyperparameters were selected to have the best response when considering the K-fold cross-validation.

## Results

The results from the four different approaches (ConvNet-LSTM and LSTM with and without data augmentation) are presented in the Table 2.

**Table 2.** Accuracy (Mean (SD)) on the validation and test set for the different model considered.

|  | K-fold cross validation<br>(K = 4) | Test |
| --- | --- | --- |
| ConvNet-LSTM + Data augmentation | 0.892 (0.010) | 0.893 |
| ConvNet-LSTM +<br>No data augmentation | 0.850 (0.012) | 0.859 |
| LSTM + Data augmentation | 0.833 (0.016) | 0.850 |
| LSTM +<br>No data augmentation | 0.797 (0.020) | 0.814 |

Moreover, the *holdout method* with the same ConvNet-LSTM hyperparameters without data augmentation was also considered. It simply consists in splitting the complete dataset into two parts randomly selected (80% training and 20% testing) in a stratified way (the same percentage of label in each set). The *holdout method* was repeated 10 times and results showed a mean accuracy of 0.997 (0.001) with the test set.

The confused logits from the ConvNet-LSTM trained with data augmentation is presented in the Table 3. For each true label (**y**), only the confused logits (***y’***) are showed. Moreover, the complete normalized confusion matrix is also presented in the supplementary materials S1 Fig as well as a Sankey diagram created using SankeyMATIC^1^ S2 Fig.

**Table 3.**
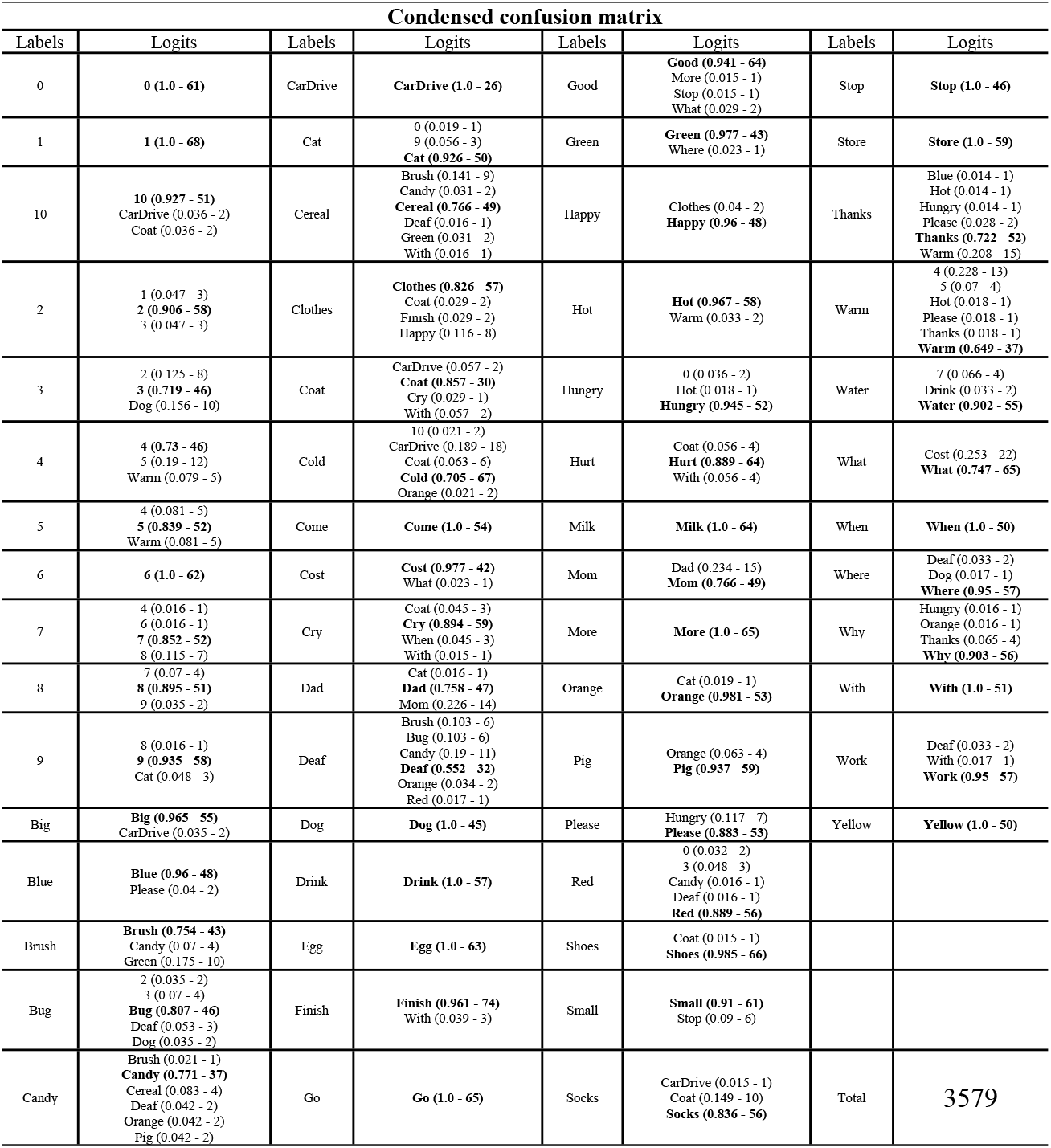
Confused logits from ConvNet-LSTM trained with data augmentation.

| Condensed confusion matrix |  |  |  |  |  |  |  |
| --- | --- | --- | --- | --- | --- | --- | --- |
| Labels | Logits | Labels | Logits | Labels | Logits | Labels | Logits |
| 0 | **0 (1.0 - 61)** | CarDrive | **CarDrive (1.0 - 26)** | Good | **Good (0.941 - 64)**<br>More (0.015 - 1)<br>Stop (0.015 - 1)<br>What (0.029 - 2) | Stop | **Stop (1.0 - 46)** |
| 1 | **1 (1.0 - 68)** | Cat | 0 (0.019 - 1)<br>9 (0.056 - 3)<br>**Cat (0.926 - 50)** | Green | **Green (0.977 - 43)**<br>Where (0.023 - 1) | Store | **Store (1.0 - 59)** |
| 10 | **10 (0.927 - 51)**<br>CarDrive (0.036 - 2)<br>Coat (0.036 - 2) | Cereal | Brush (0.141 - 9)<br>Candy (0.031 - 2)<br>**Cereal (0.766 - 49)**<br>Deaf (0.016 - 1)<br>Green (0.031 - 2)<br>With (0.016 - 1) | Happy | Clothes (0.04 - 2)<br>**Happy (0.96 - 48)** | Thanks | Blue (0.014 - 1)<br>Hot (0.014 - 1)<br>Hungry (0.014 - 1)<br>Please (0.028 - 2)<br>**Thanks (0.722 - 52)**<br>Warm (0.208 - 15) |
| 2 | 1 (0.047 - 3)<br>**2 (0.906 - 58)**<br>3 (0.047 - 3) | Clothes | **Clothes (0.826 - 57)**<br>Coat (0.029 - 2)<br>Finish (0.029 - 2)<br>Happy (0.116 - 8) | Hot | **Hot (0.967 - 58)**<br>Warm (0.033 - 2) | Warm | 4 (0.228 - 13)<br>5 (0.07 - 4)<br>Hot (0.018 - 1)<br>Please (0.018 - 1)<br>Thanks (0.018 - 1)<br>**Warm (0.649 - 37)** |
| 3 | 2 (0.125 - 8)<br>**3 (0.719 - 46)**<br>Dog (0.156 - 10) | Coat | CarDrive (0.057 - 2)<br>**Coat (0.857 - 30)**<br>Cry (0.029 - 1)<br>With (0.057 - 2) | Hungry | 0 (0.036 - 2)<br>Hot (0.018 - 1)<br>**Hungry (0.945 - 52)** | Water | 7 (0.066 - 4)<br>Drink (0.033 - 2)<br>**Water (0.902 - 55)** |
| 4 | **4 (0.73 - 46)**<br>5 (0.19 - 12)<br>Warm (0.079 - 5) | Cold | 10 (0.021 - 2)<br>CarDrive (0.189 - 18)<br>Coat (0.063 - 6)<br>**Cold (0.705 - 67)**<br>Orange (0.021 - 2) | Hurt | Coat (0.056 - 4)<br>**Hurt (0.889 - 64)**<br>With (0.056 - 4) | What | Cost (0.253 - 22)<br>**What (0.747 - 65)** |
| 5 | 4 (0.081 - 5)<br>**5 (0.839 - 52)**<br>Warm (0.081 - 5) | Come | **Come (1.0 - 54)** | Milk | **Milk (1.0 - 64)** | When | **When (1.0 - 50)** |
| 6 | **6 (1.0 - 62)** | Cost | **Cost (0.977 - 42)**<br>What (0.023 - 1) | Mom | Dad (0.234 - 15)<br>**Mom (0.766 - 49)** | Where | Deaf (0.033 - 2)<br>Dog (0.017 - 1)<br>**Where (0.95 - 57)** |
| 7 | 4 (0.016 - 1)<br>6 (0.016 - 1)<br>**7 (0.852 - 52)**<br>8 (0.115 - 7) | Cry | Coat (0.045 - 3)<br>**Cry (0.894 - 59)**<br>When (0.045 - 3)<br>With (0.015 - 1) | More | **More (1.0 - 65)** | Why | Hungry (0.016 - 1)<br>Orange (0.016 - 1)<br>Thanks (0.065 - 4)<br>**Why (0.903 - 56)** |
| 8 | 7 (0.07 - 4)<br>**8 (0.895 - 51)**<br>9 (0.035 - 2) | Dad | Cat (0.016 - 1)<br>**Dad (0.758 - 47)**<br>Mom (0.226 - 14) | Orange | Cat (0.019 - 1)<br>**Orange (0.981 - 53)** | With | **With (1.0 - 51)** |
| 9 | 8 (0.016 - 1)<br>**9 (0.935 - 58)**<br>Cat (0.048 - 3) | Deaf | Brush (0.103 - 6)<br>Bug (0.103 - 6)<br>Candy (0.19 - 11)<br>**Deaf (0.552 - 32)**<br>Orange (0.034 - 2)<br>Red (0.017 - 1) | Pig | Orange (0.063 - 4)<br>**Pig (0.937 - 59)** | Work | Deaf (0.033 - 2)<br>With (0.017 - 1)<br>**Work (0.95 - 57)** |
| Big | **Big (0.965 - 55)**<br>CarDrive (0.035 - 2) | Dog | **Dog (1.0 - 45)** | Please | Hungry (0.117 - 7)<br>**Please (0.883 - 53)** | Yellow | **Yellow (1.0 - 50)** |
| Blue | **Blue (0.96 - 48)**<br>Please (0.04 - 2) | Drink | **Drink (1.0 - 57)** | Red | 0 (0.032 - 2)<br>3 (0.048 - 3)<br>Candy (0.016 - 1)<br>Deaf (0.016 - 1)<br>**Red (0.889 - 56)** |  |  |
| Brush | **Brush (0.754 - 43)**<br>Candy (0.07 - 4)<br>Green (0.175 - 10) | Egg | **Egg (1.0 - 63)** | Shoes | Coat (0.015 - 1)<br>**Shoes (0.985 - 66)** |  |  |
| Bug | 2 (0.035 - 2)<br>3 (0.07 - 4)<br>**Bug (0.807 - 46)**<br>Deaf (0.053 - 3)<br>Dog (0.035 - 2) | Finish | **Finish (0.961 - 74)**<br>With (0.039 - 3) | Small | **Small (0.91 - 61)**<br>Stop (0.09 - 6) |  |  |
| Candy | Brush (0.021 - 1)<br>**Candy (0.771 - 37)**<br>Cereal (0.083 - 4)<br>Deaf (0.042 - 2)<br>Orange (0.042 - 2)<br>Pig (0.042 - 2) | Go | **Go (1.0 - 65)** | Socks | CarDrive (0.015 - 1)<br>Coat (0.149 - 10)<br>**Socks (0.836 - 56)** | Total | 3579 |

<sup>1</sup> http://sankeymatic.com/

For each true label of the test set composed of 3579 labels, the percentage and the absolute number of predicted logits are presented in parenthesis.

## Discussion

The aim of this study was to classify 60 signs of American Sign Language (ASL) using a deep ConvNet-LSTM with a forearm, hand and finger kinematics models from joint position data provided by the LeapMotion. The raw data from the LeapMotion for the 25 subjects is freely provided with this study (http://dx.doi.org/10.17632/8yyp7gbg6z.1). Moreover, a robust user independent K-fold cross-validation was used to create the model. This step was composed of four cycles (K = 4) with 15 subjects for training and 5 subjects for validation without overlap. It was followed by a user independent test phase with 5 subjects used to apply the model in a “real-world case” to present the classification results. Moreover, the ConvNet-LSTM was compared with a recurrent neural network composed of the LSTM cell from Zaremba, Sutskever and Vinyals (2014) to demonstrate the improvement of a combined approach.

Results showed that the ConvNet-LSTM with data augmentation showed the highest accuracy with 89.3% of the signs in the test set that were correctly estimated. The effect of data augmentation on this model showed an improvement of about 3.4% (Table 2). Regarding the LSTM, the accuracy was 85.0% and 81.4% with and without data augmentation respectively (Table 2). This difference is about the same magnitude as that of the ConvNet-LSTM demonstrating the importance of using data augmentation technique to help the classifier to generalize the human behaviour (Um et al. 2017). As presented in the study of Ordóñez and Roggen (2016), the ConvNet-LSTM outperformed the ConvNet used alone whereas in our study it outperformed the LSTM demonstrating the importance of using both approaches for HAR. Moreover, the strong point of the convolutional layer is that it could allow flexible data fusion from different sensors and data. Indeed, the 1D kernel on the columns of data considered in this study can be changed to capture inter-vector/sensor component if needed. Nevertheless, using a 1D kernel in this study provided the best results. As also pointed out by Ordóñez and Roggen (2016), using max pooling at each convolutional layers provided poorer results. Moreover, the test time was about 9.3 (0.5)s. and 36.5 (1.7)s. for the ConvNet-LSTM and LTSM respectively providing another advantage.

Regarding the *holdout method* (used only for demonstration purpose), the accuracy on the validation set was at 99.7 (0.1) %. This result showed how easy it is to gain a very high accuracy, but this method has to be avoided since the ConvNet-LSTM has already learn all the subject’s behaviour and the results would remain specific to these subjects. Such high accuracy does not necessarily mean that the neural network will provide such results with new subjects. Moreover, others ConvNet-LSTM hyperparameters that provided lower accuracy with the user independent approach also provided accuracy around 99% with the *holdout method*. Such case could lead to create a model with non-adapted hyperparameters.

It remains difficult to compare our results with those of previously reported studies since they used different validation and/or test methods, but we believe that the user independent K-fold cross-validation approach is a relevant way to prove the reliability of a model.

The Sankey diagram (Appendix B) shows for each true label y the wrong logits **y’** that has been identified in an easy-to-read manner. As presented, the words *Cold, Three, Four, Warm, Thanks, Cereal, Deaf, Mom, Dad* were the most difficult for the ConvNet-LSTM to classify (Table 3). For example, *Three* was confused with *Two* and *Four* with *Five*. Indeed, it was sometimes difficult for the LeapMotion sensor to differentiate the numbers of extended fingers. Moreover, *Cereal* and *Warm* are movement relatively close to the trunk or the head and this may have disturbed the LeapMotion on the correct hand position. In addition, *Cold* required only moving the forearms with the fists closed that may provide too limited information for the neural network given the model here used. Finally, *Mom* and *Dad* required the same hand movement to be performed except that the position in front of the head is different with the thumb position at the chin and at the forehead respectively. Despite this, *Dad* was confused with *Mom* for 22.6% in the test set showing the capability of the ConvNet-LSTM to differentiate slight difference in the global dynamics of the movement.

The main limitation of this study is that subjects who were new to sign language were recruited. Indeed, beginners may have greater variability when performing movements and future work should be considered with experts to check if there would be an improvement in the classification. In addition, a static neural network has been used and future work will have to be considered with a dynamic neural network that allows adapting the length of the input sequence coupled with an automatic signal detection method (Redmon et al. 2016). Nevertheless, a static neural network may still be considered with an automatic segmentation method and then interpolate the signal. Future study should consider another model that will be fed by the output of the ConvNet-LSTM with the purpose of predicting the most probable word based on the previous words/sentence to help the classification from a given context. Finally, sign language recognition may be improved with information provided by a camera or IMU sensors to gain information about the hand position relative to the body.

## Conclusion

This study was the first to combined approach with Convolutional Neural Network and a Recurrent Neural Network with Long-Short Term Memory cells (ConvNet-LSTM) for sign language recognition. Moreover, a robust user independent K-fold cross-validation and test phase was provided. This contrast previous work, where the validation and/or the test phase were not user independent or lack of information was provided. There are several possibilities for future work to improve these results, such as the use of experts in sign language, dynamic neural network combined with an automatic segmentation technique and additional data from a camera or IMU sensors.

## Supporting Information

**S1 Fig. 1 Normalized confusion matrix**

**S2 Fig. 2 Sankey diagram**

## Footnotes

1 http://sankeymatic.com/

